# *In vivo* multi-dimensional information-keeping in *Halobacterium salinarum*

**DOI:** 10.1101/2020.02.14.949925

**Authors:** J. Davis, A. Bisson-Filho, D. Kadyrov, T. M. De Kort, M. T. Biamonte, M. Thattai, S. Thutupalli, G. M. Church

## Abstract

Shortage of raw materials needed to manufacture components for silicon-based digital memory storage has led to a search for alternatives, including systems for storing texts, images, movies and other forms of information in DNA. Use of DNA as a medium for storage of 3-D information has also been investigated. However, two problems have yet to be addressed: first, storage of 3-D information in DNA has used objects and coding schemes which require large volumes of data; second, the medium used for DNA information-keeping has been inconsistent with qualities needed for long-term data storage. Here, we address these problems. First, we created *in vivo* DNA-encoded digital archives holding precise specifications for 3- and 4-dimensional figures with unprecedented efficiency. Second, we have demonstrated more robust and longer-lasting information-carriers than earlier repositories for DNA-based data archives by inserting digital information into the halophile, *Halobacterium salinarum*, an extremophilic archaeon. We then embedded Information-keeping halophiles into crystalline mineral salts in which similar organisms have been shown to persist *in stasis* for hundreds of millions of years. We propose that digital information archives composed in 3 or more dimensions may be inserted into halophilic organisms and preserved intact for indefinite periods of time.

## Introduction

### Digital data storage in DNA

Manufacture of silicon-based physical structures for digital information-keeping is dependent on shrinking global supplies of high-purity quartz sands. Sources of these raw materials are expected to be exhausted in less than 20 years (1). Increasing demand for data storage has therefore led to a search for suitable alternatives, including digital information storage in DNA. The capacity of DNA to hold digital archives has been under development since the 1980s (2) and a growing community of researchers have pursued this goal (3-18). To date, forms of DNA-encoded information include images, text, voice, music, movies and a computer operating system (13). At present, DNA can store at least 10^18^ bytes per mm^3^, 6 orders of magnitude greater information density than the densest data storage medium currently available (1, 19). A method to store cell lineage data across all 10^10^ cells of a mouse, illustrating the power of recording data in DNA *in vivo*, has recently been reported (18). Several examples of 3-Dimensional (3-D) information-keeping in DNA have also been demonstrated.

### 3-D encoded DNA

Crystallographer Nadrian Seeman and colleagues pioneered a form of DNA encoding often overlooked by researchers interested in DNA information storage. Seeman’s technique, later called “DNA origami,” relies on the base-pairing properties of DNA’s four nucleotides to encode 3-D information into DNA (20). In following years, Paul Rothemund’s lab refined DNA origami techniques using 7kb strands of DNA from the genome of the M13 virus which are coded to fold into desired shapes with hundreds of smaller complementary ‘staple’ strands (21-22).

More recently, researchers at ETH Zurich and a collaborating Israeli scientist have converted stereolithographic code describing the 3-D figure of a rabbit into 12kb DNA oligonucleotides (23). This 3-D encoded DNA was subsequently encapsulated in silica microbeads (24) and then mixed with thermoplastic 3-D printing materials used to print unstructured triangulated 3-D copies of the described object (rabbit). Published reports refer to this version of 3-D encoded DNA as “DNA-of-Things” (23).

While highly efficient methods have been reported for encoding DNA with digital information (25), “DNA origami” and “DNA-of-things” 3-D DNA encoding methods have been comparatively inefficient, based on *in vitro* post-processing of relatively large coded molecules (>7 kb DNA) to yield a range of complex surfaces and simple 3-D objects (20-23).

Genes prescribing 3-D structure and function are long-standing biological precedents for 3-D translations of 2-D information. Leaf curvature, for instance, is an example of 3-D outcomes encoded in 2-D patterns of growth (26). Technical advantages of ideally encoded 3-D objects include the ability to acquire, display and manipulate an unlimited number of 2-dimensional (2-D) planes; usefulness for condensing information in graphs, equations, and complex mathematical models, and for representations of architecture, engineering, anatomy, topology, cartography (geography, geology, demographics, and population maps) where individual 3-D models replace multiple 2-D models (one for each view). 3-D shapes as a form of communication and information transfer may also be more readily interpreted by people from any area of the world or from a different era, compatible with the goals of long term data storage.

### Limitations of DNA-encoded information carriers

Most efforts to use DNA as a data storage medium have involved inserting data either into “naked,” molecular DNA or into DNA contained in laboratory strains of *E. coli*. Between 1999 and 2012, all attempts at recording digital information into DNA were recorded *in vivo* (27).

After 2012, a shift to *in vitro* DNA archives was based on the notion that *in vivo* DNA data storage is unlikely to become a viable alternative for mainstream digital data storage. Two reasons are given for this: first, that while bacterial cells are smaller than most other known cells, and approximately 5 orders of magnitude smaller than microchips currently in use for data storage, *in vivo* DNA data storage has lower overall information capacity owing to the comparatively large size of bacterial cells when compared with the molecular scale of DNA; and second, that there has been a tendency to avoid highly complex tasks of creating stable modifications and/or additions to natural DNA within living cells (27).

Proponents of *in vitro* molecular DNA data storage claim that “naked” DNA digital archives could remain intact for hundreds to thousands of years when kept at reasonable temperatures and isolated from light and humidity (27). Longer term survivability of *in vitro* DNA would also require disposition in sealed, sterile environments shielded from radiation sources. If DNA is left unprotected, it is likely to be digested by enzymes commonly present in natural environments and promptly consumed by microorganisms (28). DNA is an inherently unstable substance, subject to damage by metabolic and hydrolytic processes including oxidative damage, depurination, depyrimidination, and cytosine deamination (29-33). DNA can also accumulate damage from environmental factors including UV light, ionizing radiation, exposure to genotoxic materials, and freeze-thaw cycles that can cause mechanical shearing (29, 32-36). Maintenance of environments stabilizing naked DNA is therefore essential for reducing information loss.

The process of making stable modifications and additions to natural DNA has always been part of the promise of molecular biology. *In vivo* DNA digital archives could take advantage of cellular machinery to protect and repair DNA (37-39) and to be conveniently and economically reproduced with little or no human intervention. As is the case with the DNA recovered from the past, the most secure and long-lasting way to store information in DNA may be to have it carried by an organism. Distribution and population numbers of cells could help to compensate for both lower information density and incidental data-parity errors in individual cells when compared with *in vitro* DNA information stores.

Nevertheless, laboratory domesticated strains such as of *E. coli* are particularly vulnerable. Bacterial strains that have become “workhorses” of molecular biology are – by both accident and intention – unlikely to survive outside of the laboratory (40). Despite these inherent weaknesses, negligent introduction of modified laboratory organisms into the environment may have the potential to damage natural ecosystems. Physical containment of laboratory organisms has therefore become augmented with biocontainment efforts that focus on engineered biological safeguards (41-42) to prevent recombinant organisms from surviving outside of controlled laboratory environments.

These vulnerabilities are inconsistent with qualities desired for DNA data storage. In addition to high information density, new digital storage structures will have to be robust enough to withstand environmental conditions for long periods of time and demand little at-rest energy cost (27).

### Halobacterium salinarum

Extremophilic organisms may be more resilient and versatile information-keepers than either naked DNA or laboratory strains of *E. coli*. Here, the example of *Halobacterium salinarum* (*Hsal*), a halophilic archaeon, is taken as representative of this abundant and diverse group.

*Hsal* is not found in fresh water environments. Osmotic pressure will cause the cells of most halophilic organisms (including *Hsal*) to burst when immersed in fresh water, but there are many more saline environments in which halophilic organisms can thrive than there are fresh water environments to threaten them. 97.5% of water on Earth is salt water. Just 2.5% is fresh, and only 0.3% of Earth’s fresh water is in liquid form on the surface (43-45). In addition to the ability of Hsal to survive in the broadest possible range of available aqueous environments, it can also survive an impressive range of environmental extremes.

*Hsal* is known to be extremely radiation resistant with reports of its chromosomes having been fragmented and then reassembled after very high radiation dosage (∼5 kGy gamma irradiation) (46-47). In relative terms, 90% of *E. coli* is killed with an exposure of 0.27 kGy at +5 degrees (48), which is an almost 18-fold lower dose than *Hsal’*s known level of tolerance.

*Hsal* is polyploid, with each cell harboring approximately 25 copies of each chromosome (49). While their chromosomes may be initially shattered into many fragments, complete chromosomes are reconstituted by making use of overlapping fragments in what is likely to be a form of recombinational repair aided by DNA single-stranded binding protein (47).

*Hsal* is also known to survive oxidative stress, routine exposure to high UV, thermal extremes, and desiccation (45-46). It has been found to survive high vacuum, microgravity, radiation and temperatures that characterize the near-Earth space environment (50-54). Studies suggest that the hypersaline environment in which *Halobacterium* thrives may be a factor for its resistance to desiccation, radiation and high vacuum (50).

There is also abundant evidence to indicate that organisms sharing phenotypic and genotypic similarities with *Hsal* may persist within crystals of primary mineral salts and other evaporites over periods of geologic time (55-68). In 2006, Vreeland et al. sequenced DNA of organisms discovered in ancient salt deposits from between 121 and 419 million years ago (the oldest DNA on record) and although six segments of that DNA had never been seen before in science, *Hsal* was shown to be a close genetic relative (67-68). Earlier discoveries had been made of halophiles embedded in fossil salts, but were called into question because of the high degree of similarity that appears to exist between all of the potentially ancient microbes, their DNA, and modern halophiles (68). Thus, in addition to the resistance of *Haloarchaea* to environmental extremes mentioned above, evidence indicating extremely long-term survival in fossil mineral salts suggests that they have apparently survived Great Extinction Events that extinguished most other terrestrial species on more than one occasion.

Spontaneous mutation of DNA in cells serves evolution but is unlikely to serve the purpose of long-term storage of immutable digital information. Information-keeping in halophiles could help to solve this problem. *In stasis* disposition of halophiles in mineral crystals may render them less likely to undergo processes of mutation and DNA damage and repair that actively reproducing cells would be expected to encounter much more frequently in other natural environments. Here, we aimed to study whether *Hsal* can in fact be used for data storage across long periods of time.

## Results

### Long term survival

While experiments performed over thousands of years are obviously impractical, to investigate further the hypothesis that *Hsal* cells can survive for long periods of time when entrapped in salt crystals, we started a liquid culture until late-exponential growth (O.D. ∼1.0) stage and then split it into two identical aliquots. The first aliquot was used for genomic DNA extraction, while the second aliquot was left to evaporate within the sterile environment of a laminar hood. After 5 days, all liquid media was transformed into salt crystals with pink color tones (Figure 1A). Salt crystals were then distributed in sterile test tubes, sealed with parafilm and stored at room temperature. For the past three years, speckles of salt crystals from one of these samples were regularly tested for growth in fresh liquid media every six months. Notably, in every test performed to this point, cells reached mid-exponential growth (O.D. ∼0.5) in less than 24 hours after inoculation. Figure 1B shows *Hsal* cells entrapped between salt crystals in brine from dehydrated hypersaline media.

**Figure 1:**
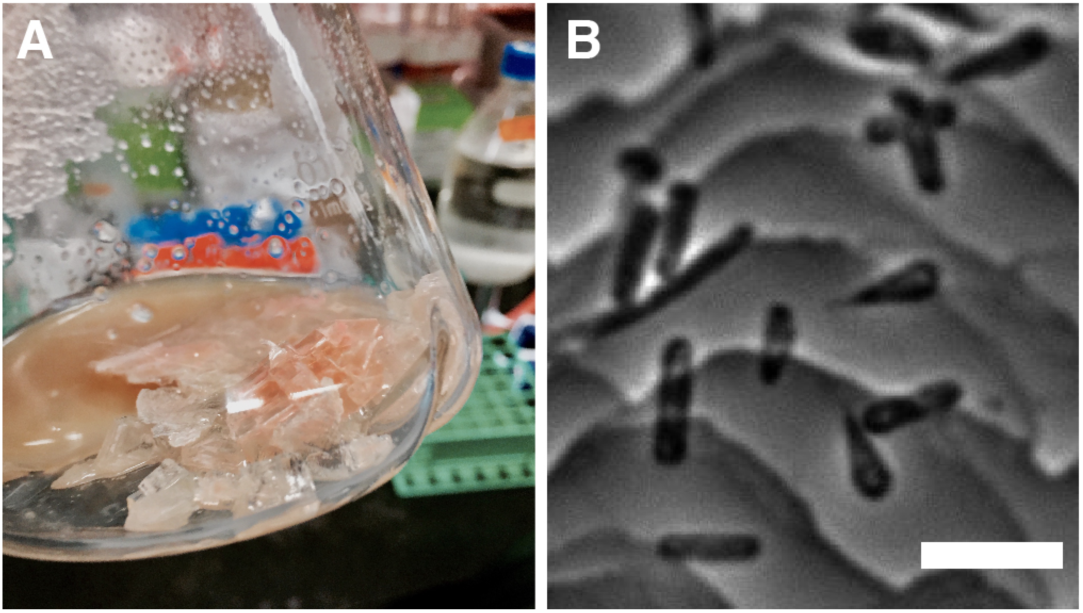
(A) Pink salts with embedded *Halobacterium salinarum* crystallized from hypersaline culture (B) Phase-contrast images of *Halobacterium salinarum* cells entrapped between salt crystals in brine from dehydrated hypersaline media. Scale bars represent 5 µm.

### 3-D information-keeping in E. coli

To bypass one of the main limitations for *in vivo* DNA storage and increase its cache capacity, we determined to expand the number of dimensions of information that a given linear DNA could carry. We utilized highly efficient vector encoding to precisely describe a 3-D double-helix object using information encoded into only 46 DNA bases (Figure 2A). From there, a traditional paper folding pattern was used to yield the 3-D double-helix (Figure 2B).

**Figure 2:**
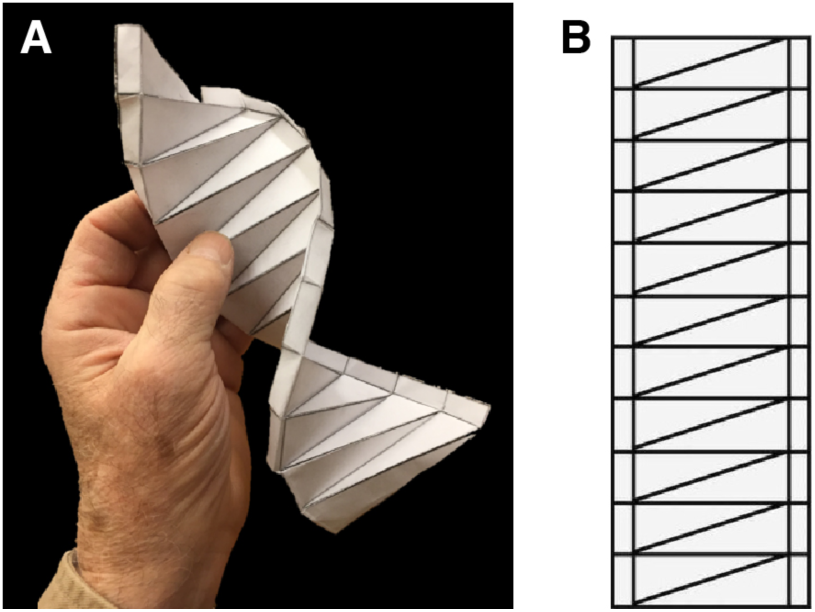
(A) 3-D origami double-helix figure (B) and corresponding origami crease pattern

Creasing patterns from origami are 2-D diagrams whose lines represent folds one must perform to transform a piece of flat paper into a 3-D origami shape (69). In origami folding using these patterns, there are two types of creases indicating the type of fold: “mountain” creases and “valley” creases. In diagrams, these two types of creases are often indicated using solid and dashed lines or using two distinct colors. In more mathematical terminology, crease patterns are planar graphs with labeled edges in which there are at most two labels (70). Our double-helix figure coding method accounts for the 2-D origami folding pattern as well as for “valley” and “mountain” creases.

The origami double-helix crease pattern was interpreted as a 94-bit lattice vector encoding 11 segments, taking advantage of the spatial periodicity of the pattern. This same method can be used to encode larger files (Supplementary Material 1). The asymptotic size of this encoding, in bits, is:

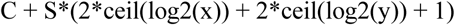

Here, ‘S’ is the number of segments (each associated with defined start and end points, with one bit encoding the folding sense, positive or negative); ‘x’ and ‘y’ give horizontal and vertical lattice resolutions; ‘C’ is a constant which depends on the maximum possible values of ‘S’, ‘x’, ‘y’. The “1” is one bit to encode positive or negative fold. This encoding must first be decoded into the corresponding 2-D origami pattern, and subsequently folded to yield the desired 3-D structure (Figure 3). DNA manipulation and *E. coli* transformations were performed according to the method of Sambrook and Russell (71). Genetic constructions were generated by conventional enzymatic restriction and ligation of inserts into vectors using competent *E. coli* MG1655 as a primary recipient. The locus was sequenced from *E. coli* and reconstructed into the 3-D double-helix using the decoded crease pattern.

**Figure 3:**
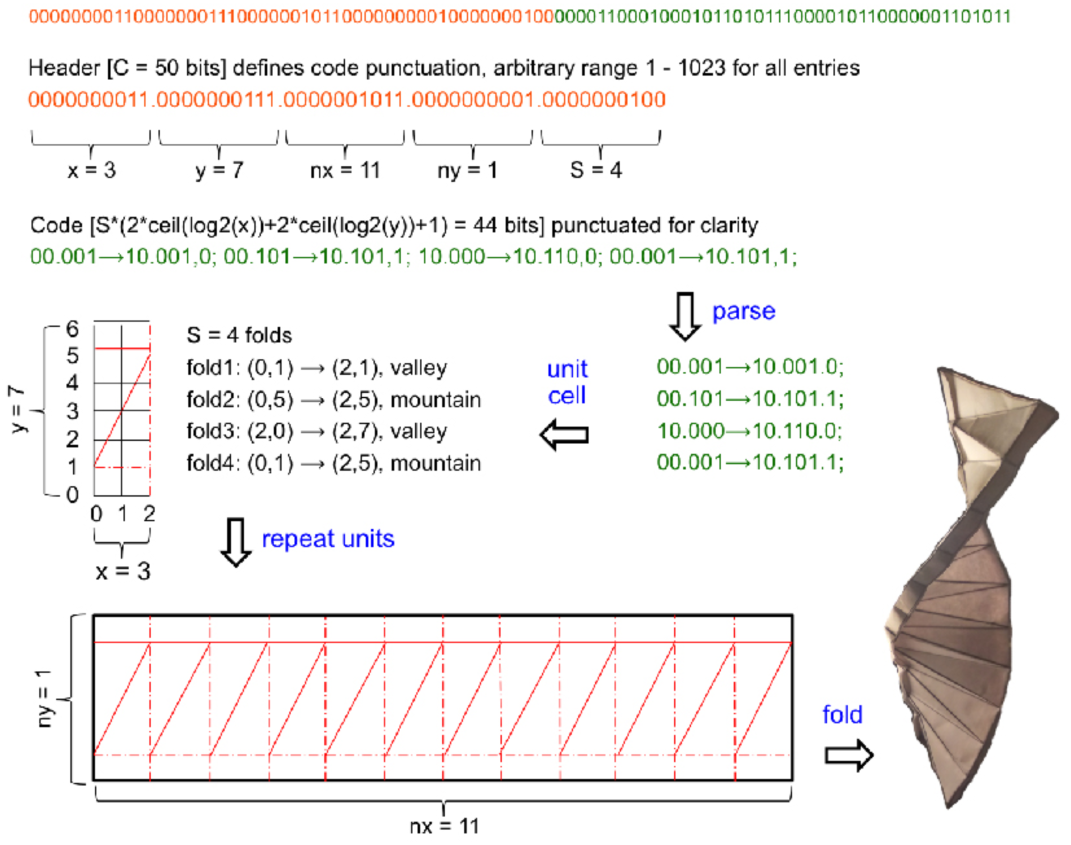
Decoding steps for converting encoded binary string into folded origami 3-D double-helix figure

### 3-D information-keeping in Halobacterium salinarum

In fall 2018, we were offered an opportunity to contribute to the immortality-themed *5*^*th*^ *Ural Industrial Biennale* (72-73), a symposium and exhibition that opened in September, 2019 in Yekaterinburg, Russia. We decided to insert 3-D digital information into *Hsal* to serve as an extremely enduring human-sourced informational archive to compliment the *Ural Biennale* theme. Content for the *Hsal* archive consisted of two 3-D objects drawn from Russian folklore about an immortal evil wizard known as “Koschei the Deathless” (Кощéй Бессмéртный)*. Based on these legends, the project was organized to insert 3-D models of a needle and an egg into the *Hsal* genome. We coded folding patterns of 3-D needle and spatial coordinates for a 3-D egg using a combination of highly efficient sparse row matrix encoding techniques (74-76) and degenerate elements inspired by “DNA Supercode” (77-80), a DNA-encoding scheme composed in the 1990s.

To generate DNA sequences with minimal lengths that encode the 3-D geometric information of the needle and egg shapes, we used techniques from both the ancient art of origami and modern data-compression algorithms. In particular, origami crease patterns were employed to compress the 3-D geometric information into 2-D diagrams. Although in general it is difficult, in the complexity-theoretic sense, to generate a 3-D origami shape from its crease pattern, the needle and egg crease patterns used in this work are simple enough that the folding instructions follow immediately from the diagrams (Figure 4) (81-82). This dimensionality reduction via crease pattern is achieved by the fact that the structure of the patterns effectively encodes the folding instructions required to transform a flat sheet of paper into a 3-D object. Typically, in origami crease patterns, two types of lines are drawn to indicate which direction the fold should be performed, called “mountain” creases and “valley” creases (Figure 5).

**Figure 4:**
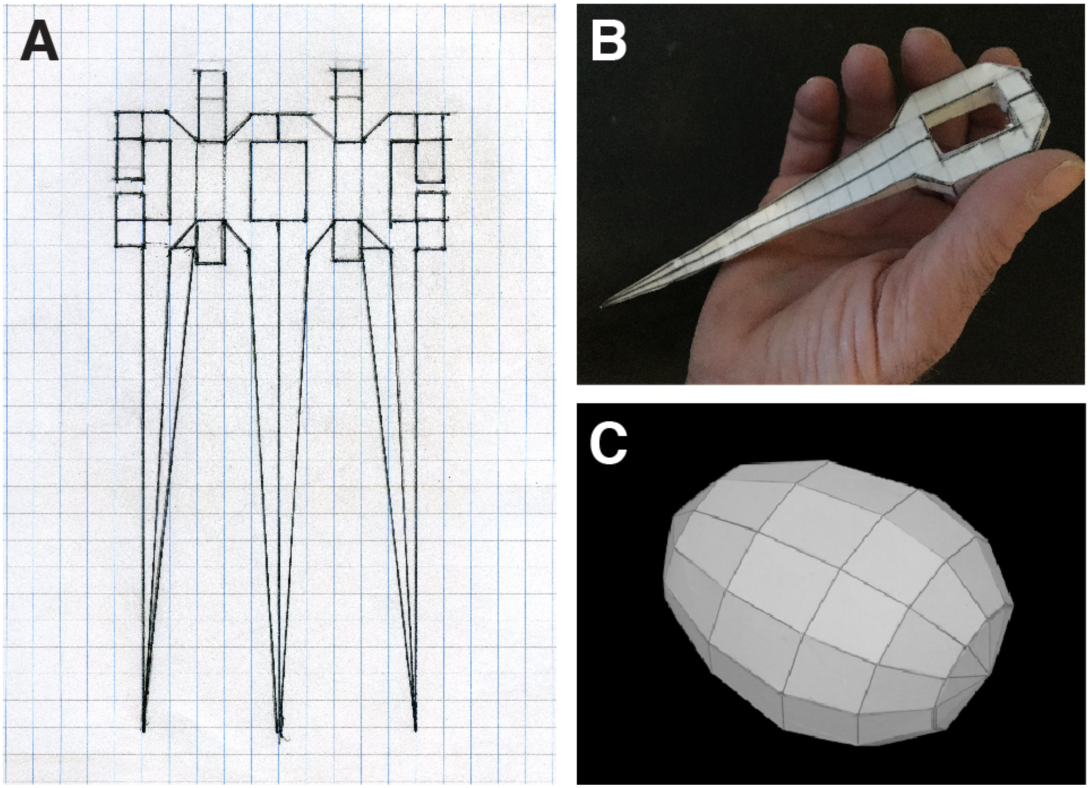
(A) Planar graph of 2-D needle crease pattern (B) 3-D folded needle, and (C) assembled 3-D egg

**Figure 5:**
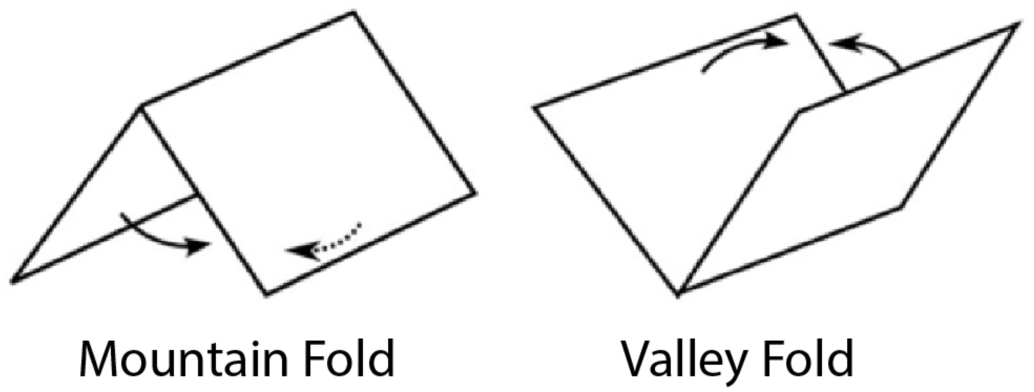
“Mountain” and “valley” origami folds

Moreover, in special cases, it is not necessary to specify types of creases needed to indicate the sequence of folds. This special situation holds for both the needle and egg diagrams employed herein. Given a target 3-D shape to be rendered, finding the crease pattern is, at the moment, still an art form. However, once a crease pattern is provided, all of the 3-D information is encoded in a 2-D diagram called a “planar graph” (Figure 4).

In this work, all creases are depicted with the same type of line; the diagram does not distinguish between mountain and valley folds. This can be done while still containing all the information to fold the 3-D object from the 2-D diagram due to a condition called “flat-foldability” in traditional origami. Flat-foldability allows for folding a shape such that it can fit between the pages of a book. Logically solving which creases are mountains and which ones are valleys is arrived at by choosing those that render a shape flat-foldable. If a given crease pattern does not distinguish mountains and valleys, the mathematically relevant features of the diagrams are the lines and the corners where the lines meet. Lines are called “edges,” and the corners where lines meet are called “vertices.” The collection of all edges and vertices together is called a graph. Here the set of vertices is denoted by V and the set of edges by E. The entire diagram is described by the graph formed from this pair of sets and is denoted G = (V,E). Locations of the vertices together with a description of which vertices are connected to each other specify our graphs. Locations of vertices are determined by superimposing a square grid over the crease pattern and recording the x and y coordinates. We chose an arbitrary labelling of vertices. Given N vertices, each were labeled with numbers from the set {1,2,3,…,N-1,N}. We imposed a square coordinate grid and chose lexicographic ordering.

To describe which vertices are connected to each other, it is standard to use a matrix – a square table of numbers – data structure. The matrix describing the connectivity of the vertices is called an “adjacency matrix.” We created adjacency matrices for each of our encoded 3-D figures (Figure 6). An adjacency matrix for an N-vertex diagram is an N by N (square) matrix whose (*i, j*) entry is one if vertex *i* is connected to vertex *j* and zero otherwise. For most crease patterns, each vertex in the diagram is not connected to every other vertex and the adjacency matrix will have many zeros. This means that for most reasonable objects, the adjacency matrix associated with the crease pattern will be a what is termed a “sparse matrix.” For visual clarity, instead of writing the numbers 0 and 1, we plot a black square if the value in the matrix is 1 and plot a white square if the value is 0.

**Figure 6:**
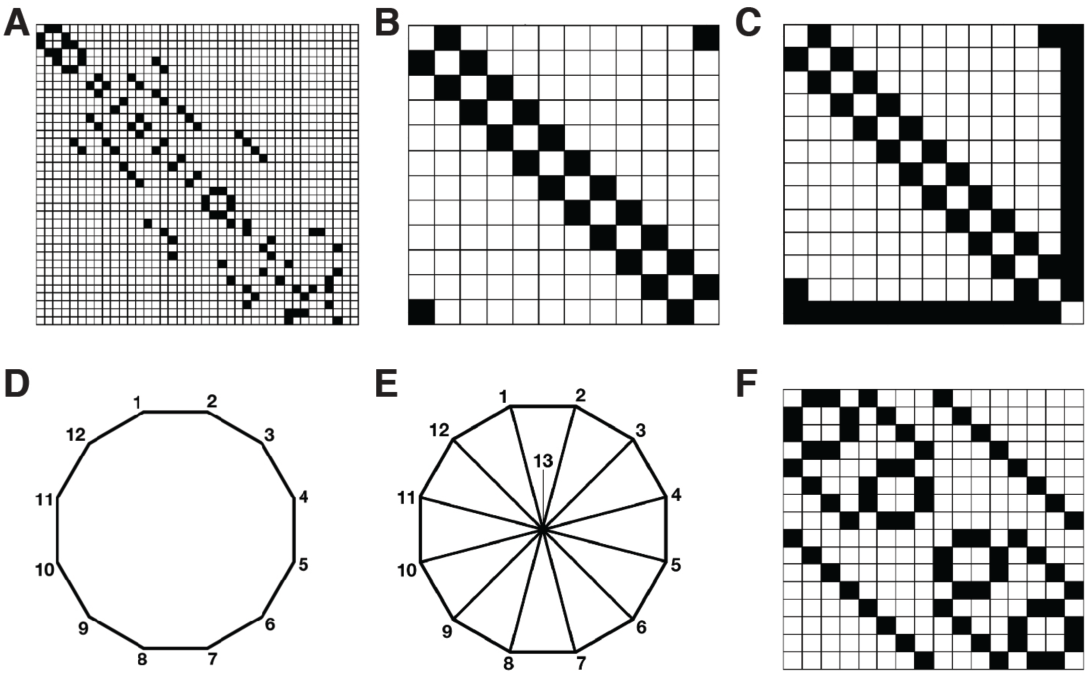
(A) Adjacency matrix for needle figure (B) Adjacency matrix for egg body figure (C) Adjacency matrix for egg end cap (with “spokes”) figure (D) Cross section of egg figure body with labeled body (dodecagon vertices) (E) Egg figure diagram for endcaps (dodecagon with spokes along with labelled vertices) (F) Adjacency matrix for the 4-D hypercube figure

Large contiguous regions of white squares represent the lack of lines between associated pairs of vertices. This is a generic feature of origami crease patterns and we use this feature to further compress data required to specify 3-D geometry. In the language of matrix theory, this feature is called “sparsity” and an adjacency matrix with a substantial fraction of its entries being zero is called a “sparse matrix.”

There exist many compression algorithms for storing sparse matrices. We chose the compressed row storage (CRS) algorithm (74-76). The algorithm compresses the information required to store the adjacency matrix by only keeping track of: 1) the values of the non-zero entries, 2) the column indices of these non-zero entries, and 3) the row indices corresponding to the first non-zero entry in each row as read from left to right. These data were converted to binary, and then to sequences of DNA bases. T**h**e resulting concatenated DNA sequence encoding needle and egg figures has a length of 930 bases (Supplementary Materials 2).

### DNA-encoding 4 dimensions

We also encoded data for a 4-dimensional hypercube or, “tesseract”** into DNA to emphasize the difference between our method – which can plausibly hold an unlimited number of dimensions – and “DNA origami” and “DNA of Things” methods, which are both *in vitro* only techniques, intensive in the size and number of prerequisite DNA molecules, and limited to 3 spatial dimensions. The 4-D hypercube was encoded via specification of the locations of its vertices together with its adjacency matrix (Figure 6). We then coded the 4-D hypercube binary data into a 224-mer DNA sequence (Supplementary Material 3).

### Assembly of Needle/Egg and Hypercube oligonucleotides

As repeats in the Needle/Egg and the Hypercube DNA sequences were predicted to cause issues during synthesis of the DNA oligos, both sequences were divided into segments, with Egg-Needle sequences being split into three segments and the Hypercube sequence into two. Each segment was augmented with BbsI recognition sites and elongated with random sequences until the complexities of the segments were sufficiently reduced (Table S1). The augmented sequences were ordered as gBlocks (Integrated DNA Technologies). The segments were subsequently assembled by Golden Gate assembly with the BbsI-HF restriction enzyme (New England Biolabs) and T7 ligase (New England Biolabs). The resulting products were amplified by PCR using Q5 High-Fidelity Master Mix (New England Biolabs) and the amplicons (Table S2) were purified with a PCR Purification kit (Qiagen). Constructs were verified using Sanger sequencing.

### Transformations with Halobacterium salinarum

The Egg-Needle and hypercube constructs were inserted into Hsal NRC-1 (ATCC700922) main chromosome at the ura3 locus by double-crossover method transformation (83), with modifications. Instead of using standard counterselection with ura3, we selected transformants by introducing a MevR resistance cassette by a 4-piece Gibson reaction (84) with the following fragments: ura3 upstream region (primers oHS273 and oHV172, amplified from Hsal NRC-1 genomic DNA); Egg-Needle fragment (obtained as described above); MevR cassette (primers oHV173 and oHS278, amplified from pNBKO7 plasmid (85-86); ura3downstream region (primers oHS279 and oHS280, amplified from Hsal NRC1-1 genomic DNA). Primer sequences are listed in Table S2. The assembled fragments were then directly transformed by the standard PEG transformation method (83). Transformants (Figure 7) were then screened by PCR and the inserted sequence was confirmed by Sanger sequencing. A locus map of the final construct is represented in Figure 7.

**Figure 7:**
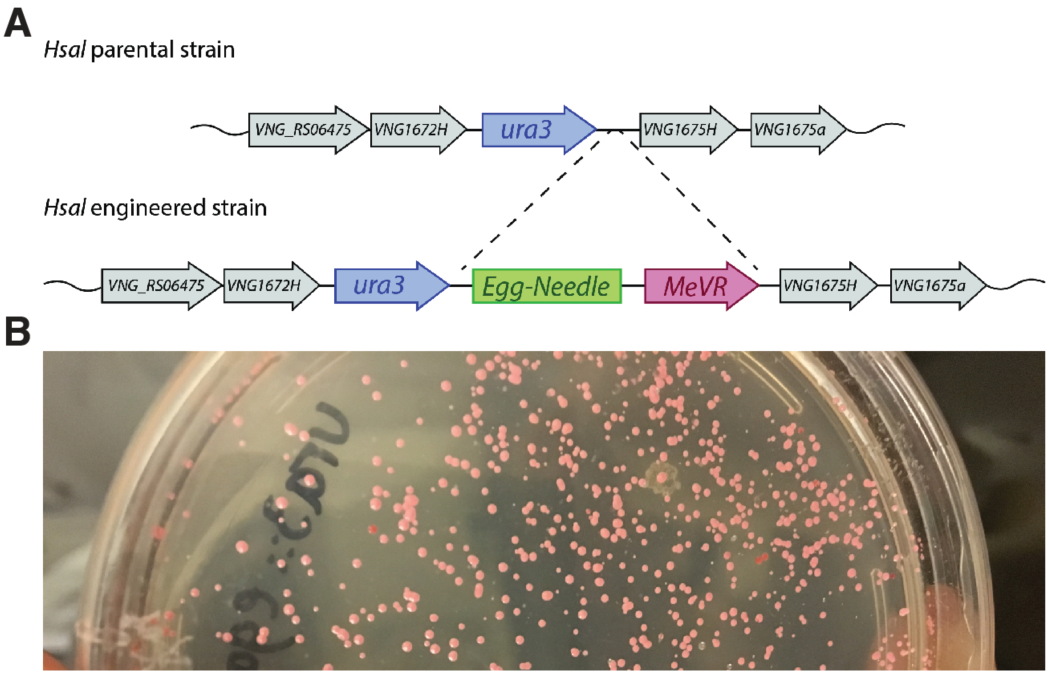
(A) Locus map of *Halobacterium salinarum* egg/needle construct.(B) Transformants holding 3-D encoded needle/egg figures.

These protocols were repeated to clone a separate *Hsal* line with DNA coding for the 4-D hypercube figure***.

### Digitally encoded Halobacterium salinarum embedded in mineral salts

Sterile brines made from various fossil mineral salts were inoculated with 3-D encoded *Hsal* and then recrystallized over a 3-day period in a series of workshops conducted at Ural Federal University Department of Experimental Biology and Biotechnology (Yekaterinburg, Russia), as part of the *Ural Biennale*.

## Discussion

### Prospects for 3-D language

While information written into sets of phonetic characters require readers to be fluent in particular languages, information written into sets of 3-D objects merely requires mathematical understandings needed to reconstruct 3-D objects from code. Some of the oldest systems of written language (e.g., cuneiform, hieroglyphics) operated by encoding information into 2-D pictures. Picture-writing systems may be useful as a first step to interpret information initially composed in the form of spoken language into more universally translatable sets of 3-D objects. Number sets could be written as sets of multifaceted polyhedra, while abstract ideas might be written as sets of contrasting, comparative, or metaphorical 3-D figures. Whether or not the nuance and poetry of language might be transcribed into 3-D figure-sets can be compared with questions about whether or not expressions in the form of painting might also be expressed in the form of sculpture.

### Legacy and burdens of transcendence

It may be short-sighted to think about digitally encoded DNA merely as a means to provide relatively near-term solutions to 21^st^ century problems in data storage. Biological information repositories can now be expected to survive for much longer than the span of time that *Homo sapiens sapiens* is likely to persist as a species.

Fossil records indicate that the average lifespan of mammalian species is roughly on the order of 1 million years (87-89). In the case of closely related hominids, Neanderthals (*Homo sapiens neanderthalensis)* survived for only 200,000-300,000 years (90-91), while *Homo erectus* survived for about 1.6 million (92-97). Scholars debate forecasts of the survivability of *Homo sapiens* for many reasons but setting aside the possibility that humanity will shortly engage in catastrophic thermonuclear exchange, predictions made over the past few decades suggest that *Homo sapiens sapiens* will survive for anywhere from ∼600 to 7.8 million years (98-100).

Humanity has inherited special awareness of mortality that may be unique among members of the animal kingdom. Presumably, this special knowledge serves as inspiration to make lasting contributions to the legacy that each individual human being may leave behind. This same knowledge can also inspire questions about what bequests our species as a whole might leave for a far deeper future.

### Cosmological time-scales

It is safe to assume that testaments written into salt-embedded microorganisms could persist over timescales that would be required for interstellar transits. The intelligibility of 3-D language might also be conveniently adapted for that purpose. But these ideas are overshadowed by the futility of communications over enormous spans that separate stars in our region of the galaxy.

Currently at a distance of 100 astronomical units (AU) and traveling outward at about 2.54 AU per year, *Pioneer 10* is one of the fastest ballistic objects ever set into motion by human beings. Traveling at about 12.13 km/s relative to the sun, the *Pioneer 10* spacecraft is approximately 271,000 AU from *Proxima Centauri* (the nearest star) (101).

If *Pioneer 10* was targeted on *Proxima Centauri* (which it is not), it would not arrive for about 106,693 years. For context, behaviorally modern humans (possessing behavioral and cognitive traits such as abstract thinking, depth of planning, uses of symbolism, and blade-making technology) are thought to have been in existence for only about 50,000 years (102) while *Homo sapiens* is thought to have existed as a species distinct from other hominids for about 200,000 years (103)

It seems safe to assume that by the time any physical message could arrive in the neighborhood of a star more remote than the sun’s nearest companions, *Homo sapiens* will either be extinct or, will have evolved into another species.

We do not know how many unique star systems, all at greater distances than *Proxima Centauri*, would need to be targeted to reach a planet where life has developed, much less a planet with sentient life having the capacity to intercept and understand any message humanity had decided to transmit. Despite sophisticated searches for signals from others, the chances that humanity will ever find itself communicating with another intelligent species somewhere else in the universe are slim to none (104).

Earth will likely outlast humanity and remain habitable to other DNA-based organisms. Whether or not future terrestrial life evolves into organisms that are remotely like human beings, if we can manage to commit the legacy of our science, culture and civilization into forms of data storage that will outlast *Homo sapiens*, we may enable communications with beings who could turn out to be the only other communicable species in the cosmos, right here on Earth.

### Swansong

The 3-D encoded “Koschei” *Hsal* has a poetic part in the form of coded figures that conjure an ancient myth about immortality. It is also first use of a medium that can reach into an indefinite future and as such, it begs the question about what serious, information-dense messages could also be projected across such extreme periods of time.

Legends about the swan who sings a last, final lament were proverbial by the 3rd century BCE (105-106) and reiterated many times in later Western poetry and art. But any ideas we may entertain about contacting beings far across space and time are complicated by another problem, one perhaps even more daunting than the distance between stars.

Understanding what intelligence is in the first place is prerequisite to reliably imagining any other intelligence or, to say it another way, “*You have to reveal yourself to yourself before you can reveal yourself to anyone else*.*”* This is what Aristotle considered to be a principal element of human tragedy (“Recognition and Reversal” in *Poetics*) (107), and yet, it is probably the most important reason we have to continue the search. Any messages we may wish to preserve for the “Other” will have to attempt to answer the questions “Is this who we are? Is this what we know?”

While “DNA origami” techniques may have useful applications, 3-D objects rendered as digitally-encoded DNA are more efficiently described as strictly mathematical constructs rather than as exotic, secondary molecular conformations.

Extremophiles can be more robust and long-lasting information-carriers than earlier repositories for DNA-based digital archives. Indeed, techniques currently exist that can be used to transfer digital information into the DNA of *Halobacterium salinarum*. Furthermore, useful applications of digital archives encoded into extremophile DNA are not limited to routine data storage systems. Other plausible applications include interstellar messaging and legacy terrestrial, lunar and planetary archives.

## Acknowledgments

The authors are indebted to art-and-science curator, Olga Vad, whose initial offering brought a large part of this work into the realm of possibility, and to Yashas Shetty, whose enthusiasm brought several of us together for the first time, and whose insights into alternative methods for encoding 3-D objects helped to start the ball rolling. We are grateful to Irina Kiselyova, head of Department of Experimental Biology and Biotechnology at Ural Federal University, Russia for the opportunity to host a series of our experiments with extremophilic archaea; and to Ido Bachelet, for his advice and support. We also acknowledge Gabriel Filsinger, Eswar Iyer, Srividya Chandramouli, Erkin Kuru, and Henry Lee for time spent reviewing the manuscript.

## Footnotes

* Koschei is said to be “deathless” because he is supposed to have hidden his evil soul in the tip of a needle which he then concealed in an egg, then in a duck, then in a rabbit, and so on, and then to have buried these concentric concealments in a chest that was buried under a tree on a mythical oceanic island (Buyan) that would appear and disappear with alternating tides.

** A tesseract is the 4-dimensional analog of the cube. A tesseract is a 4D hypercube.

*** The Needle/Egg-encoded DNA and the 4-D hypercube-encoded DNA were transformed with separate respective *Hsal* cell lines and confirmed using Sanger sequencing.

## References

(1) Zhirnov, V., Zadegan, R. M., Sandhu, G. S., Church, G. M. & Hughes, W. L. Nucleic acid memory. Nature Materials. 15, 366–370 (2016).

(2) Davis, J. Microvenus. Art Journal. 55, 70–74 (Spring 1996). [Art Journal 55 (1996) This entire issue (Ellen Levy, ed.) was devoted to Art and the Genetic Code, representing an early example of an art journal addressing issues of art and genetics]

(3) C. T. Clelland, V. Risca and C. Bancroft. Hiding messages in DNA microdots. Nature 399, 533–534 (10 June 1999)

(4) Bancroft, C. Long-term storage of information in DNA, Science 293, 1763–1765 (10 August 2001)

(5) Wong, P.C., Wong, K., Foote, H. Organic data memory using the DNA approach. Communications of the Association for Computing Machinery (ACM) 46, 95–98 (March 2003)

(6) Arita, M., Ohashi, Y. Secret signatures inside genomic DNA. Biotechnology Progress 20 1605–1607 (Sept.-Oct. 2004)

(7) Yachie, N., Sekiyama, K., Sugahara, J., Ohashi, Y., Tomita, M., Alignment-based approach for durable data storage into living organisms. Biotechnology Progress 23, 501–505 (25 Jan. 2007)

(8) Portney, N. G., Wu, Y., Quezada, L. K., Lonardi, S., Ozkan, M. Length-based encoding of binary data in DNA. Langmuir 24, 1613–1616 (04 Mar. 2008)

(9) Ailenberg, M. Rotstein, O. D. An improved Huffman coding method for archiving text, images, and music characters in DNA. Biotechniques 47, 747–754 (Sept. 2009)

(10) Church, G. M., Gao, Y., Kosuri, S. Next generation digital information storage in DNA. Science. 337, 1628 (28 Sept 2012; Epub 16 Aug 2012)

(11) Goldman, N., Bertone, P., Chen, S., Dessimoz, C., Leproust, E. M., Sipos, B. and Birney, E. Towards practical, high-capacity, low maintenance information storage in synthesized DNA. Nature 494, 77–80, (07 Feb. 2013)

(12) Bornholt, J., Lopez, R., Carmean, D., Ceze, L., Seelig, G., Strauss, K. A DNA-based archival storage system. Proceedings, 21st International Conference on Architectural Support for Programming Languages and Operating Systems (ASPLOS.; Published by ACM –Association for Computing Machinery; https://www.microsoft.com/enus/research/publication/dna-based-archival-storage-system (April 2016)

(13) Erlich, Y. & Zielinski, D. DNA Fountain enables a robust and efficient storage architecture. Science 355, 950–954 (03 Mar. 2017).

(14) Zhirnov, V., Zadegan, R. M., Sandhu, G. S., Church, G. M. & Hughes, W. L. Nucleic acid memory. Nature Materials 15, 366–370 (Apr. 2016).

(15) Shipman, S., Nivala, J., Macklis, J.D., Church G. M. CRISPR-Cas encoding of a digital movie into the genomes of a population of living bacteria. Nature. 547, 345–349 (20 July 2017; Epub 12 July 2017)

(16) Choi, Y., Ryu, T., Lee, A. C., Choi, H., Lee, H., Park, J., Song, S., Kim, S., Kim, H., Park, W., Kwon, S. High Information capacity DNA-based data storage with augmented encoding characteristics using degenerate bases. Nature Scientific Reports 9, (29 April 2019)

(17) Lee, H. H., Kalhor, R., Naveen, G., Bolot, J., Church, G. M. Termina-tor-free template-independent enzymatic DNA synthesis for digital information storage Nature Communications 10, (2019) [https://www.nature.com/articles/s41467-019-10258-1]

(18) Kalhor, R., Kalhor, K., Mejia, L., Leeper, K., Graveline, A., Mali, P., Church, G. M. Developmental barcoding of whole mouse via homing CRISPR Science 361, (Aug. 2018)

(19) Rutten, M. G. T. A., Vaandrager, F. W., Elemans, J. A. A. W. & Nolte, R. J. M. Encoding information into polymers. Nature Reviews Chemistry 2, 365–381 (2018).

(20) Chen, J. & Seeman, N. C. Synthesis from DNA of a molecule with the connectivity of a cube. Nature 350, 631–633 (18 April 1991)

(21) Rothemund, P. W. K. Folding DNA to create nanoscale shapes and patterns. Nature 440, 297–302 (16 March 2006)

(22) Rothemund, P. W. K. & Andersen, E. S. Nanotechnology: the importance of being modular. Nature 485, 584–585 (31 May 2012)

(23) Koch, J., Gantenbein, S., Masania, K., Wendelin, J. S., Erlich, Y., Grass, R. N. A DNA-of things storage architecture to create materials with embedded memory. Nature Biotechnology 38, 39–43 (Jan. 2020; Epub 09 Dec. 2019)

(24) Stark, W. J. Robust chemical preservation of digital information in silica with error-correction codes. Angewandte Chemie (International ed. English) 54, 2552–2555 (04 Feb. 2015)

(25) Choi, Y., Ryu, T., Lee, A. C., Choi, H., Lee, H., Park, J., Song, S., Kim, S., Kim, H., Park, W., Kwon, S. High information capacity DNA-based data storage with augmented encoding characteristics using degenerate bases. Nature Scientific Reports 9, (29 April 2019)

(26) Nath, U., Crawford, B. C. W., Carpenter, R., Coen, E. Genetic control of surface curvature Science 28, 1404–1407 (28 Feb., 2003)

(27) Ceze, L., Nivala, J., Strauss, K. Molecular digital data storage using DNA. Nature Reviews Genetics 20, 456–466 (Aug. 2019)

(28) Clark, D. P., Pazdernik N. J. Transformation, uptake of naked DNA. In Molecular Biology Second Edition/Academic Cell Update) *Academic Press* (Elsevier) 641–646 (2013)

(29) Lindahl, T. Instability and decay of the primary structure of DNA Nature. 362, 709–715. (April 1993).

(30) De Bont R, van Larebeke N. Endogenous DNA damage in humans: a review of quantitative data”. Mutagenesis. 19, 169–85 (May 2004)

(31) Lindahl T, Nyberg B. Rate of depurination of native deoxyribonucleic acid. Biochemistry 11, 3610–3618 (Sept. 1972).

(32) Shikama, K. Effect of freezing and thawing on the stability of double helix of DNA. Nature 207, 529–530 (July 1965)

(33) Röder, B., Frühwirth, K., Vogl, C., Wagner, M., Rossmanith, P. Impact of long-term storage on stability of standard DNA for nucleic acid-based methods Journal of Clinical Microbiology 48, 4260–4262 (Nov. 2010)

(34) Wondergem, J. (ed.) Radiation biology: a handbook for teachers and students. Vienna: International Vienna Atomic Energy Agency pp. 26, 72 (Mar. 2010)

(35) Häder D-P, Sinha RP. Solar ultraviolet radiation-induced DNA damage in aquatic organisms: potential environmental impact. Mutation Research. 571, 2005; 221–233

(36) Rastogi, R. P., Richa Kumar, A., Tyagi, M. B., Sinha, R. P. Molecular Mechanisms of ultraviolet radiation-Induced DNA damage and repair. Journal of Nucleic Acids 2010, (Dec. 2010)

(37) Wigly, D. B. Bacterial DNA repair: recent insights into the mechanism of RecBCD, AddAB and AdnAB. Nature Reviews Microbiology 11, 9–13 (Dec. 2013)

(38) Kunkel TA, Erie DA (2005). DNA mismatch repair. Annual Review of Biochemistry. 74: 681–710 (Jan. 2013).

(39) Yi, C., He, C. DNA repair by reversal of DNA damage. Cold Spring Harbor Perspectives in Biology. 5 (Jan. 2013)

(40) Fux, C. A., Shirtliff, M., Stoodley, P., Costerton, J. W. Can laboratory reference strains mirror “real world” pathogenesis? Trends in Microbiology 13, 58–32 (01 Feb. 2005)

(41) Mandell, D., Lajoie, M., Mee, M., Takeuchi, R., Kuznetsov, G., Norville, J. E., Gregg, C. J., Stoddard, B. L., Church, G. M. Biocontainment of genetically modified organisms by synthetic protein design. Nature 518, 55–60 (05 Feb. 2015; Epub 21 Jan. 2015)

(42) Castanon, O., Smith, C. J., Khoshakhlagh, P., Ferreira, R., Guell, M., Said, K., Ramazan, Y., Dysart, M., Wang, S., Thompson, D., Myllykallio, H., Church, G. M. CRISPR-mediated biocontainment https://t.co/qRJRp6M6Vr#bioRxiv (04 Feb. 2020)

(43) Shiklomanov, L. A. World fresh water resources In: Gleick P. H. (ed.), Water in Crisis: A Guide to the World’s Fresh Water Resources, New York: Oxford University Press 13–24 (1993)

(44) Where is Earth’s water? United States Geological Survey https://web.archive.org/web/20131214091601/http://ga.water.usgs.gov/edu/earthwherewater.html

(45) Eakins, B. W., Sharman, G. F. volumes of the world’s oceans from ETOPO1 Boulder: NOAA National Geophysical Data Center (2010) [ETOPO1 is a 1 arc-minute global relief model of Earth’s surface that integrates land topography and ocean bathymetry]

(46) https://science.nasa.gov/science-news/science-at-nasa/2004/10sep_radmicrobe/

(47) DeVeaux, L. C., Müller, J. A., Smith, J., Petrisko, J., Wells, D. P., DasSarma, S. Extremely Radiation-resistant mutants of a halophilic archaeon with increased single-stranded DNA-binding protein (RPA) gene expression. Radiation Research 168, 507–514 (1 Oct. 2007)

(48) Thayer, D. W., Boyd, G. Elimination of Escherichia coli 0157:H7 in meats by gamma irradiation. Applied Environmental Microbiology 59, 1030–1034 (Apr. 1993)

(49) Hildenbrand, C., Stock, T., Lange, C., Rother, M., Soppa, J. Genome copy numbers and gene conversion in methanogenic archaea. Journal of Bacteriology 18, 734–743 (Nov. 2010)

(50) Weider, G., Leuko, S., Stan-Lotter, H. Survival and growth of Halobacterium sp. NRC-1 following incubation at −15°C, freezing or freeze-drying, and the protective effect of cations. Proceedings of the Third European Workshop on Exo-Astrobiology, 18 - 20, Madrid, Spain. Ed.: Harris, R. A., Ouwehand, L. ESA SP-545 311-312 (November 2003)

(51) Kotteman, M., Kish, A., Iloanusi, C., Bjork, S., DiRuggiero, J. Physiological responses of the halophilic archaeon Halobacterium sp. Strain NRC1 to desiccation and gamma irradiation. Extremophiles 9, (June 2005; Epub 21 Apr. 2005)

(52) Robinson, J. L., Pyzyna, B., Atrasz, R. G., Henderson C. A., Morrill, K. L., Burd, A. M., DeSoucy, E., Fogleman III, R. E., Naylor, J. B., Steele, S. M., Elliott, D. R., Leyva, K. J., Shand, R. F. Growth kinetics of extremely halophilic *archaea* (family *Halobacteriaceae*) as revealed by Arrhenius plots. Journal of Bacteriology. 187, 923–929 (Feb. 2005)

(53) Cocker, J. A., DasSarma, P., Kumar, J., Müller, J. A., DasSarma, S. Transcriptional profiling of the model archaeon Halobacterium sp. NRC-1: responses to changes in salinity and temperature Saline Systems 3:6; http://www.salinesystems.org/content/3/1/6 (25 July 2007)

(54) Robinson, C. K., Webb, K., Amardeep, K., Jaruga, P., Dizdaroglu, M., Baliga, N. S., Place, A., DiRuggiero, J. A major role for nonenzymatic antioxidant processes in the radioresistance of Halobacterium salinarum. Journal of Bacteriology 193, 1653–1662 (Apr. 2011; Epub 28 Jan. 2011)

(55) Reiser, R., Tasch, P. Investigation of the viability of osmophile bacteria of great geological age. Transactions of the Kansas Academy of Science. 63, 31–34 (Jan. 1960)

(56) Dombrowski, H. Bacteria from Paleozoic salt deposits. Annals of the New York Academy of Sciences. 108, 453–460. (29 June 1963)

(57) McGenity, T. J., Gemmell, R. T., Grant, W. D., Stan-Lotter, H Origins of halophilic organisms in ancient salt deposits. Environmental Microbiology 2, 243–250 (June 2000)

(58) Gruber C., Legat, A., Pfaffenhuemer, M., Radax, C., Weidler, G., Busse, H. J., Stan-Lotter, H., Halobacterium noricense sp. nov., an archaeal isolate from a bore core of an alpine Permian salt deposit, classification of Halobacterium sp. NRC-1 as a strain of H. salinarum and emended description of H. salinarum Extremophiles 8, 431–439 (08 Dec. 2004; Epub 30 Jul 2004) 2004.

(59) Schubert, B. A., Lowenstein, T. K., Timofeeff, M. N., Parker, M. A. Halophilic archaea cultured from ancient halite, Death Valley, California. Environmental Microbiology 12, 440–454 (Feb. 2010)

(60) Lowenstein, T. K., Schubert, B. A., Timofeeff, M. N. Microbial communities in fluid inclusions and long-term survival in halite. GSA Today (The Geological Society of America) 21, 4–9 (Jan. 2011)

(61) Sankaranarayanan, K., Lowenstein, T. K., Timofeeff, M. N., Schubert, B. A., Lum, J. K. Characterization of ancient DNA supports long-term survival of haloarchaea. Astrobiology 14, 553–560 (July 2014)

(62) Schinteie, R., Brocks, J. J. Evidence for ancient halophiles? Testing biomarker syngeneity of evaporates from Neoproterozoic and Cambrian strata. Organic Geochemistry 72, 46–58 (July 2014)

(63) Jaakkola ST, Zerulla K, Guo Q, Liu Y, Ma H, et al. Halophilic archaea cultivated from surface sterilized middle-late Eocene rock salt are polyploid. PLoS ONE 9(10): (22 Oct. 2014)

(64) Stan-Lotter, H., Fendrihan, S. Halophilic archaea: life with desiccation, radiation and oligotrophy over geologic time. Life (Basel) 5, 1487–1496 (Sept. 2015; Epub 28 July 2015)

(65) Jaakkola, S. T., Ravantti, J. J., Oksanen, H. M., Bamford, D. H. Buried alive: microbes from ancient halite. Trends in Microbiology 24, 148–160 (February 2016)

(66) Megaw, J., Kelly S. A., Thompson, T. P., Skvortsov, T., Gilmore, B. F. Profiling the microbial community of a Triassic halite deposit in Northern Ireland: an environment with significant potential for biodiscovery. FEMS Microbiology Letters 366, (28 November 2019)

(67) Vreeland, H; Rosenzweig, W D; Lowenstein, T; Satterfield, C; Ventosa, A (December 2006). Fatty acid and DNA analyses of Permian bacteria isolated from ancient salt crystals reveal differences with their modern relatives. Extremophiles 10, 71–78 (Dec. 2006)

(68) Park, J. S., Vreeland, R. H., Cho, B. C., Lowenstein, T. K., Timofeeff, M. N., Rosenzweig, W. D. Haloarchaeal diversity in 23, 121 and 419 MYA salts. Geobiology 7, 515–523 (2009)

(69) Akitaya, H. A., Mitani, J., Kanamori, Y., Fukui, Y. Generating folding sequences from crease patterns of flat-foldable origami. ACM SIGGRAPH 2013 Posters. 1-1 (July 2013).

(70) Bender E. A., Edward A., Williamson, S. G. Lists, Decisions and Graphs. San Diego: S. Gill Williamson/University of California at San Diego (Dec. 2010).

(71) Sambrook J, Russell, D. W. Molecular cloning: a laboratory manual CSHL Press (2001)

(72) https://fifth.uralbiennale.ru/en/2019/08/28/library/

(73) https://fifth.uralbiennale.ru/en/program/workshop/

(74) Bai, Zhaojun, Demmel, J., Dongarra, J., Ruhe, A., van der Vorst, H., (eds.) Templates for the solution of algebraic eigenvalue problems: a practical guide. Philadelphia: Society for Industrial and Applied Mathematics (2000)

(75) Buluç, Aydin, Fineman, J. T., Frigo, M., Gilbert, J. R., Leiserson, C. E. Parallel sparse matrix-vector and matrix-transpose-vector multiplication using compressed spare blocks (PDF) Association for Computer Machinery (ACM) Symposium on Parallelism in Algorithms and Architectures (2009)

(76) https://en.wikipedia.org/wiki/Sparse_matrix

(77) http://90.146.8.18/en/archiv_files/20001/E2000_249.pdf

(78) Davis, J. Romance, supercodes and the Milky Way DNA. In: Stocker G., Schopf (eds.) *Ars Electronica 2000 Catalog: Next Sex*, 217-235, Vienna, Austria: Springer-Verlag (2000)

(79) Davis, J., Boyd., D., O’Reilly, H., Wieczorek, M. Art and genetics In: Cooper, D. N., (ed.) Nature Encyclopedia of the Human Genome, London; New York: Macmillan Publishers Ltd, Nature Publishing Group: (2003)

(80) Davis, J., Boyd., D., O’Reilly, H., Wieczorek, M. Art and genetics In: Encyclopedia of Life Sciences (ELS) ed. (2006; Epub 15 Sept. 2006)

(81) Lang, R. J. Origami design secrets: mathematical methods for an ancient art. AK Peters/CRC Press, (2011).

(82) Arkin, E. M., Bender, M. A., Demaine, E. D., Demaine, M. L., Mitchell, J. S., Sethia, S., & Skiena, S. S. When can you fold a map? Computational Geometry, 29, 23–46 (2004).

(83) Peck, R. F., DasSarma, S., Krebs, M. P. Homologous gene knockout in the archaeon Halobacterium salinarum with ura3 as a counterselectable marker. Molecular Microbiology 35, 667–676 (Feb. 2000)

(84) Gibson, D. G., Young, L., Chuang, R. Y., Venter, J. C., Hutchison C. A., 3^rd^, Smith H. O. Enzymatic assembly of DNA molecules up to several hundred kilobases. Nature Methods 6, 343–345 (May 2009; Epub 12 Apr. 2009)

(85) Busch, C. R. DNA mismatch repair and response to oxidative stress in the extremely halophilic archaeon Halobacterium sp. strain NRC-1 [Dissertation submitted to the Faculty of the Graduate School of the University of Maryland, College Park, in partial fulfillment of the requirements for the degree of Doctor of Philosophy 2008] [first reference to pNBKO7]

(86) Schmidt, D., Wilson, M. D., Spyrou, C., Brown, G. D., Hadfield, J., Odom, D. T. ChIP-seq: using high-throughput sequencing to discover protein-DNA interactions Methods 48, 240–248 (July 2009)

(87) Wilson, E. O., Half Earth: our planet’s fight for life (First edition.). New York: Liveright Publishing Corporation, a division of W.W. Norton & Company. (07 March 2016) [“History Redefined” Chapter 16: “The average span across all groups combined appears to be (very roughly) a million years.”]

(88) Lawton, J. H., May, R. M., Extinction Rates Oxford: Oxford University Press. (01 Jan. 1995).

(89) The current mass extinction [https://www.pbs.org/wgbh/evolution/library/03/2/l_032_04.html]

(90) Monnier, G. Neanderthal behavior. Nature Education Knowledge 3, 11(2012)

(91) Alper, J., Rethinking Neanderthals Smithsonian (June 2003)

(92) https://www.livescience.com/41048-facts-about-homo-erectus.html

(93) Ferring, R.; Oms, O.; Agusti, J.; Berna, F.; Nioradze, M.; Shelia, T.; Tappen, M.; Vekua, A.; Zhvania, D.; Lordkipanidze, D. Earliest human occupations at Dmanisi (Georgian Caucasus) dated to 1,85-1.78 Ma Proceedings of the National Academy of Sciences. 108, 10432–10436. (2011).

(94) Garcia, T., Féraud, G., Falguères, C., de Lumley, H., Perrenoud, C., & Lordkipanidze, D. Earliest human remains in Eurasia: New 40Ar/39Ar dating of the Dmanisi hominid-bearing levels, Georgia. Quaternary Geochronology, 5, 443–451. (2010)

(95) Van Arsdale, A. P. Homo erectus - A bigger, smarter, faster hominin lineage. Nature Education Knowledge 4, (2013)

(96) Antón, S. Natural history of *Homo erectus*. American Journal of Physical Anthropology 122, 126–170 (2003).

(97) https://www.nhm.ac.uk/discover/homo-erectus-our-ancient-ancestor.html

(98) Benétreau-Dupin, Y., (ed.) Doomsday argument San Francisco State University (2019); https://philpapers.org/browse/doomsday-argument

(99) Leslie, J. A., The end of the world: the science and ethics of human extinction New York: Routledge (28 Mar. 1996)

(100) Hawking, S., Tencent WE Summit 2017 keynote: https://www.youtube.com/watch?v=U-hcSLya0_w [ca. 2017 Stephen Hawking predicted extinction of Homo sapiens in 583 years.]

(101) Peat, C. (Heavens-Above GmbH) Spacecraft escaping the Solar System. (Archived at) Wayback Machine (27 March 2007): https://web.archive.org/web/20070427184732/http://www.heavensabove.com/solar-escape.asp

(102) Petragalia, M. D., Korisettar, R. (ed.) Early human behaviour in global context: the rise and diversity of the lower Paleolithic record London: Routledge (Dec. 1998)

(103) Hammond, A. S.; Royer, D. F.; Fleagle, J. G. The Omo-Kibish I pelvis. Journal of Human Evolution. 108, 199–219 (Jul 2017).

(104) Scharf, C., Even if the Milky Way is teeming with spacefaring aliens, we should not be surprised that Earth Remains unvisited Scientific American 322, 32–39 (Jan. 2020)

(105) Aesop The Complete Fables (Transl. Temple, R., Temple, O.) Penguin Classics New York: Penguin Putnam Inc. p. 127 (1998).

(106) Arnott, W G. Swan Songs. Greece & Rome. 24, 149–153. (Oct. 1977)

(107) Aristotle Poetics (Transl. Kenny, A.) Oxford World Classics Oxford: Oxford University Press. chapters 10-12 (2013)

