## supplemental data for "*In vivo* multi-dimensional information-keeping in *Halobacterium salinarum*"

**This document includes:**

Supplemental Materials 1 to 3
Tables S1 and S2

**<u>Supplementary Material 1: DNA-encoding method for origami double-helix</u>**

We use a straightforward encoding of an origami folding pattern, in which the only sources of compression are (i) rational approximations for coordinates of fold endpoints; (ii) periodic repeats, if any. Further compression is possible by making use of correlations arising from folding, e.g. constraints of angles around vertices, etc.

In this scheme, the double-helix is encoded in the following 94-bit string:

000000001100000000111000000101100000000000100000000100000001100010001011010111000001
0110000001101011

This string is decoded as described in **Figure 3**.

The binary data are further compiled as DNA bases incremented by molecular weight where:

C = 00, T = 01, A = 10, G = 11

46-mer DNA oligonucleotide encoding 3-D double helix figure:

5'- CCCCACCCTACCCAACCCCCCCCCCGCCGCTCTTAAGATTACCTAAG - 3'

Two respective copies of the 3-D encoded oligo were inserted into the pMSC1 cosmid:

Plasmid (cosmid): pMSC1

Restriction sites:

EcoR1 and BamH1

and

Xho1 and Pst1

EcoR1: GAATTC
Spacer: GTGAC
Final: GTGACGAATTC

BamH1: GGATCC
Spacer: CAA
Final: GGATCCCAA

66-mer 3DNA oligo_1 (EcoR1-BamH1):

5' - GTGACGAATTCCCCCACCCTACCCAACCCCCCCCCGCCGCTCTTAAGATTACCTAAG GGATCCCAA - 3'

Xho1: CTCGAG
Spacer: TGATG
Final: TGATGCTCGAG

Pst1: CTGCAG
Spacer: ACGGT
Final: CTGCAGACGGT

68-mer 3DNA oligo_2 (Xho1-Pst1):

5' - TGATGCTCGAGCCCCACCCTACCCAACCCCCCCCCGCCGCTCTTAAGATTACCTAAG CTGCAGACGGT - 3' *

* Expressing this pattern as a conventional digital picture file would require tens of thousands of bits.

**Supplementary Material 2: DNA-encoding method for 3-D needle and egg figures**

**Needle figure**

The data structure of needle line set is similar to compressed row storage of sparse matrices, where:

G = points and lines
V = points
E = lines
binstr = binary string

START = 100101 = CGC
SPACE = 100110 = ATT
END = 100111 = ATG

(* s – space character *)

Assuming unit increment in the x-coordinate direction in iteration over needle graph, needle contains y coordinates in lexicographical order until the 's' character is reached. Once the 's' character is reached, increment x

Read coordinates in lexicographical order at constant x-coordinate until reaching 's' character; when 's' character is reached increment x-coordinate by one unit.

NEEDLE_LENGTH = 101000 = AAC

needleV: =

{(0,4),(0,5),(2,4),(2,5),(3,4),(3,5),(4,1),(4,2),(4,3),(4,6),(4,7),(5,1),(5,2),(5,3),(5,4),(5,5),(5,6),(5,7),(6,1),(6,2),(7,1),(7,2),(8,1),(8,2),(8,3),(8,4),(8.5),(8,6),(8,7),(9,1),(9,2),(9,3),(9,4),(9,5),(9,6),(10,3),(10,7)}

needleE: =

{2,3,s,1,4,s,1,4,s,2,3,s,6,10,s,5,7,11,s,6,13,s,9,14,s,5,11,s,6,10,12,18,s,11,23,s,7,14,s,8,13,s,16,26,s,9,15,s,10,18,s,11,17,s,20,21,s,19,22,s,20,21,23,29,s,12,22,s,30,31,33,s,32,34,s,15,27,s,21,29,s,22,28,35,s,24,31,35,s,25,36,s,24,34,s,25,33,s,29,30,31,s,27,32}

[Convention used for needle line set -> compressed row storage format]

**Conversion of needle figure points and lines sets to binary strings**

Programming code statements (for converting coordinates to binary strings) are from the *Mathematica* programming language.

needleVbinary: =

StringJoin[StringPadLeft[Replace[Flatten[needleV],{0->"0",1->"1",2->"10",3->"11",4->"100",5->"101",6->"110",7->"111",8->"1000",9->"1001",10->"1010"},{1}],6,"0"]]

1000000000001010000100001000000100001010000110001000000110001010001000000001000
1000000100001000000110001000000110000100000111000101000000100010100001000010100
0011000101000100000101000101000101000110000101000111000110000001000110000010000
1110000001000111000010001000000000100100000000100010000000110010000000100001000000
1010010000001100010000001110010010000010010010000100010010000110010010001000001
0010001010010010001100010100000110010100001 11

NeedleEbinary: =

StringJoin[StringPadLeft[Replace[needleE],{0->”0",1->"1",2->"10",3->"11",4->"100",5->"101",6->"110",7->"111",8->"1000",9->"1001",10->"1010",11->"1011",12->"1100",13->"1101",14->"1110",15->"1111",16->"10000",17->"10001",18->"10010",19->"10011",20->"10100",21->"10101",22->"10110",23->"10111",24->"11000",25->"11001",26->"11010",27->"11011",28->"11100",29->"11101",30->"11110",31->"11111",32->"100000",33->"100001",34->"100010",35->"100011",36->"100100",s->"100101",r->"100110"},{1}],6,"0"]]

100000111001010000001000100100101000000100010010010100001000000111001010001100010
1010010100010100011100101110010100011000110110010100100100111010010100010100010
1110010100011000101000110001001010010100101101011110010100011100111010010100010
0000110110010101000000110101001010010010011111001010010100100101001010010110100
0110010101010001010110010101001101011010010101010001010101011101110110010100 11
0001011010010101111001111110000110010110000001000101001010011110110111001010101
0101110110010101011001110010001110010101100000111111000111001010110011001001001
0101100010001010010101100110000110010101110101111001111110010101101110000 0

**Needle figure binary conversion to DNA:**

Numerical base 4 (DNA) converted numerical base 2 where (based on molecular weight):

C = 00, T = 01, A = 10, G = 11

needleVdnaSeq: =

"CCCCTCCCCCTTCCACTCCCACTTCCGCTCCCGCTTCTCCCTCTCCCACTCCCGCTCC
TACTCCTGCTTCCTCTTCCACTTCCGCTTCTCCTTCTTCTTCTACTTCTGCTACCTCTA
CCACTGCCTCTGCCACACCCTCACCCACACCCGCACCTCCACCTTCACCTACACCTG
CATCCTCATCCACATCCGCATCTCCATCTTCATCTACAACCGCAACTG"

needleEdnaSeq: =

"CCACCGATTCCTCTCATTCCTCTCATTCCACCGATTCTACAAATTCTTCTGCAGATTC
TACGTATTCATCGAATTCTTCAGATTCTACAACGCTCAATTCAGTTGATTCTGCGAAT

TCACCGTATTTCCTAAATTCATCGGATTCAATCAATTCAGTCTATTTTCTTTATTTCGT
TAATTTTCTTTTTGTGTATTCGCTTAATTTGATGGACTATTACCACAATTCGGTAGAT
TTTTTGTATTTTATGCACGATTTACTGGACGATTTATATCATTTACACAATTTATACT
ATTTGTTGATGGATTTAGACC"

needleGdnaSeq: = START NEEDLE_LENGTH SPACE needleV SPACE needleE END =

"ATTAACCCCCTCCCCCTTCCACTCCCACTTCCGCTCCCGCTTCTCCCTCTCCCACTCC
CGCTCCTACTCCTGCTTCCTCTTCCACTTCCGCTTCTCCTTCTTCTTCTACTTCTGCTA
CCTCTACCACTGCCTCTGCCACACCCTCACCCACACCCGCACCTCCACCTTCACCTAC
ACCTGCATCCTCATCCACATCCGCATCTCCATCTTCATCTACAACCGCAACTGCGCCC
ACCGATTCCTCTCATTCCTCTCATTCCACCGATTCTACAAATTCTTCTGCAGATTCTA
CGTATTCATCGAATTCTTCAGATTCTACAACGCTCAATTCAGTTGATTCTGCGAATTC
ACCGTATTTCCTAAATTCATCGGATTCAATCAATTCAGTCTATTTTCTTTATTTCGTTA
ATTTTCTTTTTGTGTATTCGCTTAATTTGATGGACTATTACCACAATTCGGTAGATTTT
TTGTATTTTATGCACGATTTACTGGACGATTTATATCATTTACACAATTTATACTATT
TGTTGATGGATTTAGACCATG”

Needle figure DNA sequence is 547 base pairs long.

**Egg figure:**

EGG_LENGTH _ 1 = 101001 = AAT (1.25)
EGG_LENGTH _ 2 = 101010 = AAA (1.5)
EGG_LENGTH _ 3 = 101011 = AAG (2.25)
EGG_LENGTH _ 4 = 101100 = AGC (2.5)

Egg figure length parameters scale:

1st endcap: 1.25
1st body ring: 2.25
2nd body ring: 2.5
3rd body ring: 2.25
2nd cap endcap: 1.5

(V or E) 1-dodecagon points and lines for body of egg
(V or E) 2-dodecagon points and lines for egg endcaps

Dodecagon points are stored in row-by-row order

Y coordinates are listed

Rows separated by space characters

EggV1: =

{2,3,s,1,4,s,0,5,s,0,5,s,1,4,s,2,3} (dodec BODY points)

EggV2: =

{2,4,s,1,5,s,0,6,s,3,s,0,6,s,1,5,s,2,4} (dodec ENDCAP points)

EggE1: =

{2,3,s,1,4,s,1,5,s,2,6,s,3,7,s,4,8,s,5,9,s,6,10,s,7,11,s,8,12,s,9,12,s,10,11} (dodec BODY lines)

EggE2: =

{2,3,13,s,1,4,13,s,1,5,13,s,2,6,13,s,3,7,13,s,4,8,13,s,5,9,13,s,6,10,13,s,7,11,13,s,8,12,13,s,9,12,13,s,10,11,13} (dodec ENDCAP lines)

**Conversion of egg figure decimal coordinates to binary:**

V1eggbinary: =

StringJoin[Replace[StringPartition[StringJoin[StringPadLeft[Replace[dodecBODYpoints,{0->"0",1->"1",2->"10",3->"11",4->"100",5->"101",6->"110",7->"111",8->"1000",9->"1001",10->"1010",11->"1011",12->"1100",13->"1101",s->"100101"},{1}],6,"0"]],2],{"00"->"C","01"->"T","10"->"A","11"->"G"},{1}]]

0000100000111001010000010001001001010000000001011001010000000001011001010000010001001001010000010000011

V2eggbinary =

StringJoin[Replace[StringPartition[StringJoin[StringPadLeft[Replace[dodecENDCAPpoints,{0->"0",1->"1",2->"10",3->"11",4->"100",5->"101",6->"110",7->"111",8->"1000",9->"1001",10->"1010",11->"1011",12->"1100",13->"1101",s->"100101"},{1}],6,"0"]],2],{"00"->"C","01"->"T","10"->"A","11"->"G"},{1}]]

100001001001010000001000101100101000000000110100101000011100101000000000110100101000000100010110010100000100000100

E1eggbinary: =

StringJoin[Replace[StringPartition[StringJoin[StringPadLeft[Replace[dodecENDCAPpoints,{0->"0",1->"1",2->"10",3->"11",4->"100",5->"101",6->"110",7->"111",8->"1000",9->"1001",10->"1010",11->"1011",12->"1100",13->"1101",s->"100101"},{1}],6,"0"]],2],{"00"->"C","01"->"T","10"->"A","11"->"G"},{1}]]

100001001001010000001000101100101000000000110100101000011100101000000000110100101000000100010110010100000100000100

E2eggbinary: =

StringJoin[Replace[StringPartition[StringJoin[StringPadLeft[Replace[dodecENDCAPlines,{0->"0",1->"1",2->"10",3->"11",4->"100",5->"101",6->"110",7->"111",8->"1000",9->"1001",10->"1010",11->"1011",12->"1100",13->"1101",s->"100101"},{1}],6,"0"]],2],{"00"->"C","01"->"T","10"->"A","11"->"G"},{1}]]

10000011001101100101000001000100001101100101000001000101001101100101000010000110001101100101000011000111001101100101000100001000001101100101000101001001001101100101000110001010001101100101000111001011001101100101001000001100001101100101001001001100001101100101001010001011001101

**Egg figure binary conversion to DNA:**

V1eggDNAseq =

“CCACCGATTCCTCTCATTCCCCTTATTCCCCTTATTCCTCTCATTCCACCG"

V2eggDNAseq =

"CCACTCATTCCTCTTATTCCCCTAATTCCGATTCCCCTAATTCCTCTTATTCCACTC"

E1eggDNAseq =

"CCACCGATTCCTCTCATTCCTCTTATTCCACTAATTCCGCTGATTCTCCACATTCTTC ATATTCTACAAATTCTGCAGATTCACCGCATTCATCGCATTCAACAG"

E2eggDNAseq =

“CCACCGCGTATTCCTCTCCGTATTCCTCTTCGTATTCCACTACGTATTCCGCTGCGTA TTCTCCACCGTATTCTTCATCGTATTCTACAACGTATTCTGCAGCGTATTCACCGCCG TATTCATCGCCGTATTCAACAGCGT"

eggGdnaSeq: =

START EGG_LENGTH _ENDCAP _ 1 EGG_LENGTH _BODY _RING _ 1 EGG_LENGTH _BODY _RING _ 2 EGG_LENGTH _BODY _RING _ 3 EGG_LENGTH _ENDCAP _ 2 SPACE V1eggDNAseq SPACE E1eggDNAseq SPACE V2eggDNAseq SPACE E2eggDNAseq END =

"ATTAATAAGAGCAAGAAACCACCGATTCCTCTCATTCCCCTTATTCCCCTTATTCCT CTCATTCCACCGCGCCCACCGATTCCTCTCATTCCTCTTATTCCACTAATTCCGCTGA TTCTCCACATTCTTCATATTCTACAAATTCTGCAGATTCACCGCATTCATCGCATTCA ACAGCGCCCACTCATTCCTCTTATTCCCCTAATTCCGATTCCCCTAATTCCTCTTATTC CACTCCGCCCACCGCGTATTCCTCTCCGTATTCCTCTTCGTATTCCACTACGTATTCC

GCTGCGTATTCTCCACCGTATTCTTCATCGTATTCTACAACGTATTCTGCAGCGTATT
CACCGCCGTATTCATCGCCGTATTCAACAGCGT”

Egg figure DNA sequence is 383 base pairs long.

**930 base-pair concatenated DNA sequence encoding needle and egg figures:**

ATTAACCCCCTCCCCCTTCCACTCCCACTTCCGCTCCCGCTTCTCCCTCTCCCACTCCC
GCTCCTACTCCTGCTTCCTCTTCCACTTCCGCTTCTCCTTCTTCTTCTACTTCTGCTAC
CTCTACCACTGCCTCTGCCACACCCTCACCCACACCCGCACCTCCACCTTCACCTACA
CCTGCATCCTCATCCACATCCGCATCTCCATCTTCATCTACAACCGCAACTGCGCCCA
CCGATTCCTCTCATTCCTCTCATTCCACCGATTCTACAAATTCTTCTGCAGATTCTAC
GTATTCATCGAATTCTTCAGATTCTACAACGCTCAATTCAGTTGATTCTGCGAATTCA
CCGTATTTCCTAAATTCATCGGATTCAATCAATTCAGTCTATTTTCTTTATTTCGTTAA
TTTTCTTTTTGTGTATTCGCTTAATTTGATGGACTATTACCACAATTCGGTAGATTTTT
TGTATTTTATGCACGATTTACTGGACGATTTATATCATTTACACAATTTATACTATTT
GTTGATGGATTTAGACCATGATTATTAATAAGAGCAAGAAACCACCGATTCCTCTCA
TTCCCCTTATTCCCCTTATTCCTCTCATTCCACCGCGCCCACCGATTCCTCTCATTCCT
CTTATTCCACTAATTCCGCTGATTCTCCACATTCTTCATATTCTACAAATTCTGCAGA
TTCACCGCATTCATCGCATTCAACAGCGCCCACTCATTCCTCTTATTCCCCTAATTCC
GATTCCCCTAATTCCTCTTATTCCACTCCGCCCACCGCGTATTCCTCTCCGTATTCCTC
TTCGTATTCCACTACGTATTCCGCTGCGTATTCTCCACCGTATTCTTCATCGTATTCTA
CAACGTATTCTGCAGCGTATTCACCGCCGTATTCATCGCCGTATTCAACAGCGT

**Supplementary Material 3: 4-D hypercube-encoding into DNA**

hypercubePOINTS=StringJoin@Replace[StringPartition[StringJoin@{"0000","0001","0010","0011","0100","0101","0110","0111","1000","1001","1010","1011","1100","1101","1110","1111"},2],{"00"->"C","01"->"T","10"->"A","11"->"G"},{1}]
CCCTCACGTCTTTATGACATAAAGGCGTGAGG
StringLength["CCCTCACGTCTTTATGACATAAAGGCGTGAGG"]
32

hypercubeLINES=StringJoin@Replace[StringPartition[StringJoin@StringPadLeft[Replace[Flatten[Table[Flatten[Position[Normal[AdjacencyMatrix[HypercubeGraph[4]]][[i]],1]]-1,{i,1,Length[hypercubePOINTS]}]],{0->"0",1->"1",2->"10",3->"11",4->"100",5->"101",6->"110",7->"111",8->"1000",9->"1001",10->"1010",11->"1011",12->"1100",13->"1101",14->"1110",15->"1111"},{1}],6,"0"],2],{"00"->"C","01"->"T","10"->"A","11"->"G"},{1}]

CCTCCACTCCACCCCCCGCTTCATCCCCCGCTACAACCTCCACTGCAGCCCCTTCTAC
GCCCTCTCCTGCGTCCACTCCTGCGACCGCTTCTACGGCCCCATCAACGCCCTCACC
AGCGTCCACACCAGCGACCGCATCAACGGCTCCACCGTCGACTTCATCGCCGGCTAC
AACGCCGGCTGCAGCGTCGA

StringLength["CCTCCACTCCACCCCCCGCTTCATCCCCCGCTACAACCTCCACTGCAGC
CCCTTCTACGCCCTCTCCTGCGTCCACTCCTGCGACCGCTTCTACGGCCCCATCAACG
CCCTCACCAGCGTCCACACCAGCGACCGCATCAACGGCTCCACCGTCGACTTCATCG
CCGGCTACAACGCCGGCTGCAGCGTCGA"]
192

**Hypercube (Points and Lines) 224-mer DNA sequence**
[concatenated points with lines; no space characters]

CCCTCACGTCTTTATGACATAAAGGCGTGAGGCCTCCACTCCACCCCCCGCTTCATC
CCCCGCTACAACCTCCACTGCAGCCCCTTCTACGCCCTCTCCTGCGTCCACTCCTGCG
ACCGCTTCTACGGCCCCATCAACGCCCTCACCAGCGTCCACACCAGCGACCGCATCA
ACGGCTCCACCGTCGACTTCATCGCCGGCTACAACGCCGGCTGCAGCGTCGA

StringLength["CCCTCACGTCTTTATGACATAAAGGCGTGAGGCCTCCACTCCACCCCC
CGCTTCATCCCCCGCTACAACCTCCACTGCAGCCCCTTCTACGCCCTCTCCTGCGTCC
ACTCCTGCGACCGCTTCTACGGCCCCATCAACGCCCTCACCAGCGTCCACACCAGCG
ACCGCATCAACGGCTCCACCGTCGACTTCATCGCCGGCTACAACGCCGGCTGCAGCG
TCGA"]
224

**DNA encoding of a 4-D hypercube**
hypercubePOINTS=StringJoin@Replace[StringPartition[StringJoin@{"0000","0001","0010","0011","0100","0101","0110","0111","1000","1001","1010","1011","1100","1101","1110","1111"},2],{"00"->"C","01"->"T","10"->"A","11"->"G"},{1}]
CCCTCACGTCTTTATGACATAAAGGCGTGAGG

StringLength["CCCTCACGTCTTTATGACATAAAGGCGTGAGG"]
32

hypercubeLINES=StringJoin@Replace[StringPartition[StringJoin@StringPadLeft[Replace[Flatten[Table[Flatten[Position[Normal[AdjacencyMatrix[HypercubeGraph[4]]][[i]],1]]-1,{i,1,Length[hypercubePOINTS]}]],{0->”0",1->"1",2->"10",3->"11",4->"100",5->"101",6->"110",7->"111",8->"1000",9->"1001",10->"1010",11->"1011",12->"1100",13->"1101",14->"1110",15->"1111"},{1}],6,"0"],2],{"00"->"C","01"->"T","10"->"A","11"->"G"},{1}]

CCTCCACTCCACCCCCCGCTTCATCCCCCGCTACAACCTCCACTGCAGCCCCTTCTAC
GCCCTCTCCTGCGTCCACTCCTGCGACCGCTTCTACGGCCCCATCAACGCCCTCACC
AGCGTCCACACCAGCGACCGCATCAACGGCTCCACCGTCGACTTCATCGCCGGCTAC
AACGCCGGCTGCAGCGTCGA

StringLength["CCTCCACTCCACCCCCCGCTTCATCCCCCGCTACAACCTCCACTGCAGC
CCCTTCTACGCCCTCTCCTGCGTCCACTCCTGCGACCGCTTCTACGGCCCCATCAACG
CCCTCACCAGCGTCCACACCAGCGACCGCATCAACGGCTCCACCGTCGACTTCATCG
CCGGCTACAACGCCGGCTGCAGCGTCGA"]
192

**Hypercube Points and Lines DNA sequence**
[concatenated points with lines; no space characters]

CCCTCACGTCTTTATGACATAAAGGCGTGAGGCCTCCACTCCACCCCCCGCTTCATC
CCCCGCTACAACCTCCACTGCAGCCCCTTCTACGCCCTCTCCTGCGTCCACTCCTGCG
ACCGCTTCTACGGCCCCATCAACGCCCTCACCAGCGTCCACACCAGCGACCGCATCA
ACGGCTCCACCGTCGACTTCATCGCCGGCTACAACGCCGGCTGCAGCGTCGA

StringLength["CCCTCACGTCTTTATGACATAAAGGCGTGAGGCCTCCACTCCACCCCC
CGCTTCATCCCCCGCTACAACCTCCACTGCAGCCCCTTCTACGCCCTCTCCTGCGTCC
ACTCCTGCGACCGCTTCTACGGCCCCATCAACGCCCTCACCAGCGTCCACACCAGCG
ACCGCATCAACGGCTCCACCGTCGACTTCATCGCCGGCTACAACGCCGGCTGCAGCG
TCGA"]
224

**Table S1:** gBlock sequences. **Bold** letters indicate extensions. *Italicized* letters indicate BbsI recognition sites.

| Name | Sequence 5’-3’ |
|---|---|
| Needle-Egg1 | ATTAACCCCCTCCCCCTTCCACTCCCACTTCCGCTCCCGCTTCTCCCTCT<br>CCCACTCCCGCTC<br>CTACTCCTGCTTCCTCTTCCACTTCCGCTTCTCCTTCTT<br>CTTCTACTTCTGCTACCTCTACCAC<br>TGCCTCTGCCACACCCTCACCCACA<br>CCCGCACCTCCACCTTCACCTACACCTGCATCCTCAT<br>CCACATCCGCATCTCCATCTTCATCTACAACCGCAACTGCGCCCACCGA<br>TTCCTCTCATTCC<br>TCTCATTCCACCGATTCTACAAATTCTTCTGCAGATTCTACGTATTCATC<br>GAATTCTTCAGA<br>TTCTACAACGCTCAATTCAGTTGATTCTGCGAATTCACCGTATTTCCTA<br>ATC**GT*CTTCTG*TG GTG** |
| Needle-Egg2 | **A*GAAGAC*GA**CTAAATTCATCGGATTCAATCAATTCAGTCTATTTTCTTT<br>ATTTCGTTAATTT<br>TCTTTTTGTGTATTCGCTTAATTTGATGGACTATTACCACAATTCGGTAG<br>ATTTTTTGTATTT<br>TATGCACGATTTACTGGACGATTTATATCATTTACACAATTTATACTAT<br>TTGTTGATGGATT<br>TAGACCATGATTATTAATAAGAGCAAGAAACCACCGATTCCTCTCATTC<br>CCCTTATTCCCCT<br>TATTCCTCTCATTCCACCGCGCCCACCGATTCCTCTCATTCCTCTTATTC<br>CACTAATTCCGCT<br>GATTCTCCACATTCTTCATATTCTACAAATTCTGCAGATTCACCGCATTC<br>ATCGCATTCAAC TC**GT*CTTCTG*TGGTG** |
| Needle-Egg3 | **CACCACA*GAAGAC*GA**CAACAGCGCCCACTCATTCCTCTTATTCCCCTA<br>ATTCCGATTCCCCT<br>AATTCCTCTTATTCCACTCCGCCCACCGCGTATTCCTCTCCGTATTCCTC<br>TTCGTATTCCACTA<br>CGTATTCCGCTGCGTATTCTCCACCGTATTCTTCATCGTATTCTACAACG<br>TATTCTGCAGCGT ATTCACCGCCGTATTCATCGCCGTATTCAACAGCGT |
| Hypercube1 | CCCTCACGTCTTTATGACATAAAGGCGTGAGGCCTCCACTCCACCCCCC<br>GCTTCATCCCCCG<br>CTACAACCTCCACTGCAGCCCCTTCTACGCCCTCTCCTGCGTCCACTCC<br>TGCTC**GT*CTTCTG*T GGTG** |
| Hypercube2 | **CACCACA*GAAGAC*GA**CTGCGACCGCTTCTACGGCCCCATCAACGCCCT<br>CACCAGCGTCCA<br>CACCAGCGACCGCATCAACGGCTCCACCGTCGACTTCATCGCCGGCTA<br>CAACGCCGGCTG CAGCGTCGA**ATTAAAA** |

**Table S2:** Oligonucleotides used for PCR amplification/sequencing

| Primer name | Sequence 5'-3' |
|---|---|
| Egg-Needle Fw | ATTAACCCCCTCCCCCTTC |
| Egg-Needle Rv | ACGCTGTTGAATACGGCG |
| Hypercube Fw | CCCTCACGTCTTTATGACATAAAG |
| Hypercube Rv | TTTTAATTCGACGCTGCAGC |
| oHS273 | CTACGACGTGGCCCA |
| oHV172 | GGGGGAGGGGGTTAATGCGTCTGCTCGATCT |
| oHV173 | GCCGTATTCAACAGCGTTCGAGTCGTCCCACG |
| oHS278 | GGTATGAGCGTGACCGCATGGGAGGGGATGG |
| oHS279 | CGGTCACGCTCATACC |
| oHS280 | CAGTCGATCCGGTCG |
